# Evolution of Key Factors Influencing Performance Across Phases in Junior Short Sprints

**DOI:** 10.1101/2024.10.16.618610

**Authors:** Kyosuke Oku, Yoshihiro Kai, Hitoshi Koda, Megumi Gonno, Maki Tanaka, Tomoyuki Matsui, Yuya Watanabe, Toru Morihara, Noriyuki Kida

## Abstract

Sprint performance plays a crucial role in various sports. Short sprints vary in court size and competitive characteristics but are common in many sports. Although the relationship between age and muscle strength has been explored in short sprints, there is limited understanding of how various physical factors interact, particularly concerning differences in the acceleration phase. This study examined the relationship between sprint times at 0‒2.5 m, 2.5‒5 m, and 5‒10 m intervals and various factors in junior athletes. Results indicated that sprint times increased with age, and is correlated with muscle strength and flexibility. Partial correlation analysis showed that faster times in the 0‒2.5 m interval were associated with higher hip flexibility; in the 2.5‒5 m interval, faster times were associated with higher core flexibility; and in the 5‒10 m interval, a relationship with standing long jump performance was confirmed. Furthermore, lower fat-free body weight translated to higher performance. In the acceleration phase of 10 m, flexibility immediately after the start and the subsequent horizontal propulsive force is important factors that are strongly related to performance change in each interval. The study results illuminate the mechanism behind short sprints in junior athletes and emphasize the importance of daily training, considering the sprint distance required for each sport.

## Introduction

In various sports, it is necessary to demonstrate maximum performance within a limited space and time. Short sprints are considered crucial, particularly in sports involving locomotion. In tennis, sprints of less than 20 m have been highlighted as important (20), and predicting the time for a 5 m sprint can be performed based on four factors: age, height, lower limb explosive strength (LLES), and skill level (21). The relationship between performance and short sprints has also been reported in other sports such as handball, where differences in 30 m sprint times are based on skill level (26), and in hockey, where short sprints can serve as predictive factors for skill level (11,12). Short sprints are closely related to performance in various sports and increasing the speed of short-distance movements leads to improved performance.

During short sprints, the acceleration phase immediately after the start of the sprint is crucial. The required sprint distance varies depending on the sport; however, the acceleration phase immediately after the start is common in all sports. Previous studies have shown differences in times of 2.5 m and 5 m based on the stepping technique at the start (13), and starting with one foot down rather than setting both feet parallel at the start has been confirmed to result in faster times at 5 m and 10 m (9). Thus, the times for the initial 2.5 m, 5 m, and 10 m have been well studied, highlighting the importance of performance at these distances. Additionally, differences in the 5 m time are already evident at the 2.5 m point (13), suggesting that analysis across sections of 0‒2.5 m, 2.5‒5 m, and 5‒10 m separately may reveal important factors in each interval.

Short sprints are related to physical characteristics, such as age and body weight. A study examining 30 m sprints by age group 11‒15 years reported that as age increased, maximum speed, stride length, and ground contact time increased (27). Regarding physical factors, body weight has a negative impact on maximum speed in 30 m sprints (28). In tennis, body mass index (BMI) has been reported to affect athletic performance (15). Therefore, considering the changes in body composition with age and growth, it is important to examine the key factors in short sprints.

Physical fitness and muscle strength are important factors for short sprints. Previous studies have shown a correlation between peak velocity in vertical jumps and 10 m sprint times (25), with vertical jumps also correlating with sprint times at 10, 30, and 60 m (32). In a previous study, when participants were divided into two groups based on back squat scores, no significant time differences were observed in muscle strenght at 5 m, but individuals with stronger lower limb muscles were faster at 10 m and 20 m (7). There have also been reports of a positive correlation between half-back squats and speed at the 5 m mark (4). Thus, the numerical values of lower limb muscle strength and physical fitness tests are important in explaining short sprint performance. This study focused on lower-limb muscle strength and physical fitness tests to examine short sprints.

Maintaining physical flexibility is also important in short sprints. Previous studies have reported that the higher the flexibility of the hamstrings, the faster the times for 5 m, 10 m, and 20 m sprints (16). In this study, we examined the flexibility of the lower limbs and that of the upper body. This allowed for a detailed investigation of the functionality of short sprints involving the entire body.

We aimed to investigate the key factors that strongly influence short sprints during the junior years, such as age, body composition, physical fitness tests, muscle strength, and flexibility, at intervals of 0‒2.5 m, 2.5‒5 m, and 5‒10 m. Short sprints are crucial in various sports, especially the performance immediately after the start, which is common in many competitions. By dividing short sprints into smaller intervals and examining the changes in related factors, we can clarify detailed performance immediately after the start. Furthermore, focusing on the junior years — a period of significant age-related physical development — and comprehensively examining the relationship with body composition, muscle strength, flexibility, and motor skills adds further significance to this study. Understanding the mechanism of short sprint performance across sections of 0‒2.5 m, 2.5‒5 m, and 5‒10 m is expected to be beneficial in sports coaching and rehabilitation settings.

## Method

### Participants

This study involved 30 boys and girls aged 9‒14 years who regularly participated in competitive sports (16 boys, 14 girls, average age 11.37 ± 1.30 years). The participants specialized in the following disciplines: 8 in badminton, 10 in fencing, 5 in rowing, and 7 in climbing, with each engaging in specialized training sessions approximately twice a week. Before the experiment, the purpose and procedure were explained to the participants and written informed consent was obtained. This study was approved by the Ethics Committee of the Kyoto Institute of Technology and conducted in accordance with the Declaration of Helsinki.

### Short sprint

An overview of the measurement environment is presented in Fig. 1. All measurements were conducted indoors with the participants using indoor shoes they brought with them. Short sprints were measured using a phototube (Dashr-Blue) placed at four positions: 0 m (start point), 2.5 m, 5 m, and 10 m (finish line). The height of the phototube was set at 80 cm. Participants were asked to stand still at a position where the tip of one foot touched the starting line and then to start running from a convenient position. The starting time was set at the discretion of the participants. Each participant performed the trial twice, and the best performance was selected for analysis. Passage times at 0‒2.5 m, 2.5‒5 m, and 5‒10 m were calculated for analysis.

**Figure 1.**
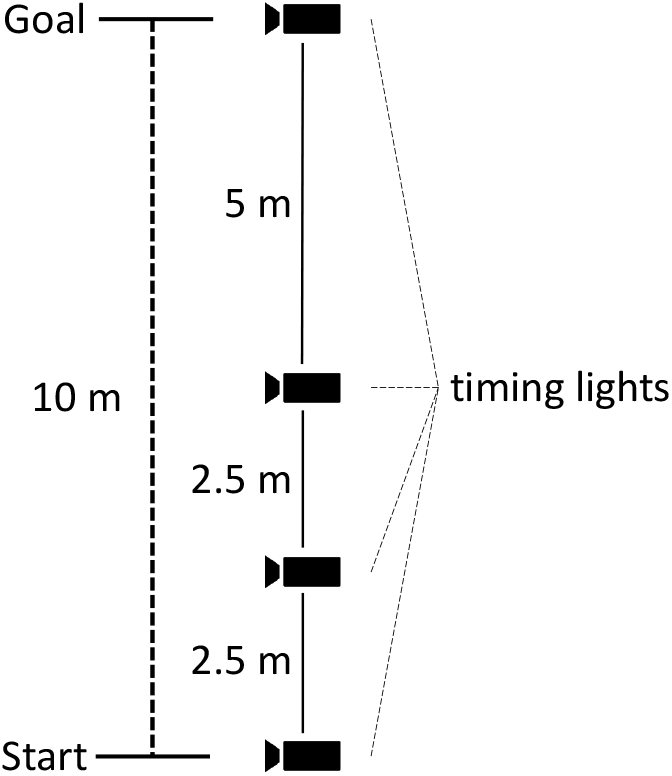
Overview diagram of short sprint measurement.

### Body composition

Body composition was measured using the InBody 570 (InBody, Inc.) for body weight, body fat mass, and fat-free mass.

### Flexibility

The measurement of joint range of motion was conducted in accordance with methods outlined in previous studies (19). Trunk rotation was performed with the pelvis fixed in a seated position. The angles between the lines connecting the left and right posterior superior iliac spines and the left and right acromion were measured. The hip flexion angle was measured with the pelvis fixed in a supine position, with the knee joint flexed naturally, and the angle between a line parallel to the trunk and the line connecting the femur (center of the greater trochanter and the lateral epicondyle of the femur) was measured. Measurements were conducted by groups of four physical therapists, with one therapist fixing the subject’s body to prevent compensatory movements, one therapist moving the subject’s body, one therapist measuring the angles and distances, and one therapist recording the data. Angles were measured in one-degree increments using a goniometer.

### Muscle strength

In the muscle strength measurements, knee extension and flexion strengths were measured. Knee extension and flexion strength were measured using Cybex Inc. in a seated position by measuring isometric and isokinetic strength during extension and flexion movements. Isokinetic strength measurements were performed at 300°/s and all participants underwent measurements on their right leg.

### Physical fitness test

Physical fitness tests included sit-and-reach, repeated side-steps, standing long jumps, sit-ups, and whole-body reaction times. Each participant was assigned an assistant to ensure safety during the fitness tests and other measurements. For the sit-and-reach test, participants sat on the ground with their backs against a wall, their legs fully extended, and were instructed to reach forward. The change in values from the initial arm extension to the post-reach position was recorded, and the best of two attempts was used (22,24). For the repeated side-step, three lines were set 1 m apart, and participants were asked to side-step back and forth for 20 s. The number of times both feet crossed each line was counted and the best result of the two attempts was used (3,33). In the standing long jump, the participants slightly widened their feet, aligned their toes with the starting line, and jumped forward simultaneously with both feet. The distance from the starting line to the back of the heels was measured, and the best result from two attempts was used (2,29,31). For the sit-ups, the participants crossed their arms over their chests, kept their knees bent at a 90° angle, and had their feet secured by an assistant. The number of times the elbows touched both thighs within a 30 s period was measured. Each participant was given only one attempt (8,18). Whole-body reaction time was measured by standing on a mat sensor and reacting as quickly as possible to visual stimuli placed at eye level by jumping slightly. The time from visual stimulus presentation to the feet leaving the mat switch was recorded (14).

### Statistical analysis

We conducted the analysis on 26 participants after excluding data from four individuals with missing values. We performed a correlation analysis between the sprint times in each short sprint interval and various variables. Subsequently, we conducted a partial correlation analysis, with age in months as the control variable.

## Result

The relationship between age in months and time in the 0‒2.5 m, 2.5‒5 m, and 5‒10 m intervals is shown in Figure 2. Negative correlations were found between age in month and time in all intervals, indicating that sprint times in each interval decreased with an increase in age (0‒2.5 m: *r* = -0.49, *p* = 0.01; 2.5‒5 m: r = -0.47, *p* = 0.02; 5‒10 m: r = -0.54, *p* = 0.004).

**Figure 2.**
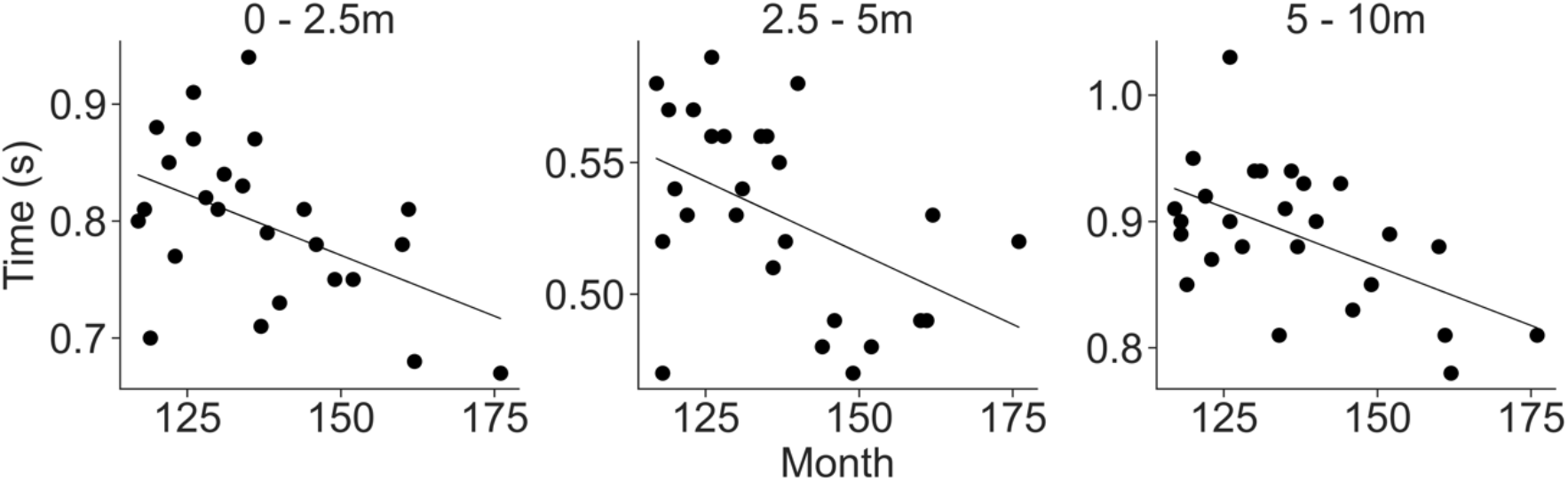
Relationship between age in month and sprint time in each interval.

We conducted a simple correlation analysis between sprint times in the 0‒2.5 m, 2.5‒5 m, and 5‒10 m intervals and various variables such as body composition, flexibility, muscle strength, and physical fitness tests (Table 1). In the 0-2.5 m interval, significant correlations were found with sprint time in the 5‒10 m interval (*r* = 0.64, *p* = 3.81×10-4), isometric knee flexion (*r* = -0.48, *p* = 0.014), and standing long jump (*r* = -0.59, *p* = 1.35×10-3). In the 2.5-5m interval, correlations were found with right-left trunk rotation (right: *r* = -0.47, *p* = 0.017; left: *r* = -0.53, *p* = 0.006), isometric knee extension (*r* = -0.45, *p* = 0.023), and standing long jump (*r* = -0.41, *p* = 0.038). In the 5‒10 m interval, significant correlations were found between sprint time (*r* = 0.64, *p* = 3.81×10-4), fat free mass (*r* = -0.42, *p* = 0.032), body fat mass (*r* = 0.45, *p* = 0.022), height (*r* = -0.45, *p* = 0.022), isometric knee flexion (*r* = -0.60, *p* = 0.001), isotonic knee flexion (*r* = -0.44, *p* = 0.023), repeated side-step (*r* = -0.39, *p* = 0.049), standing long jump (*r* = -0.72, *p* = 3.42×10-5), and whole-body reaction time (*r* = 0.40, *p* = 0.044).

**Table 1.** Correlation between sprint times in each section.

|  | 0-2.5m | 2.5-5m | 5-10m |
| --- | --- | --- | --- |
| 0-2.5 m (0.67 - 0.94 sec) | - | 0.17 | **0.64 \*\*\*** |
| 2.5-5 m (0.47 – 0.59 sec) | 0.17 | - | 0.24 |
| 5-10 m (0.78 – 1.03 sec) | **0.64 \*\*\*** | 0.24 | - |
| Age in month (117 – 176 month) | **-0.49 \*** | **-0.47 \*** | **-0.54 \*\*** |
| Height (125.9 – 170.8 cm) | -0.23 | -0.21 | **-0.45 \*** |
| Weight (22.2 – 64.9 kg) | -0.02 | 0.06 | -0.03 |
| Fat free mass (20.5 – 51.7 kg) | -0.31 | -0.17 | **-0.42 \*** |
| Body fat mass (1.1 – 28.1) | 0.34 † | 0.31 | **0.45 \*** |
| Trunk rotation (right) (42 – 81 ° ) | 0.04 | **-0.46 \*** | -0.08 |
| Trunk rotation (left) (38 – 76 ° ) | -0.17 | **-0.53 \*\*** | -0.22 |
| Hip flexion angle (right) (20 – 55 ° ) | 0.07 | 0.27 | 0.30 |
| Hip flexion angle (left) (21 – 46 ° ) | -0.25 | 0.13 | -0.02 |
| Isometric knee extension (39 – 163 kg) | -0.26 | **-0.45 \*** | -0.32 |
| Isometric knee flexion (18 – 84 kg) | **-0.48 \*** | -0.35 † | **-0.60 \*** |
| Isotonic knee extension (20 – 69 kg) | -0.28 | -0.23 | -0.34 † |
| Isotonic knee flexion (14 – 45 kg) | -0.22 | -0.37 † | **-0.44 \*** |
| Repeated side-step (35 – 64 times) | -0.37 † | -0.35 † | **-0.39 \*** |
| Sit-ups (23 – 38 times) | -0.37 † | -0.28 | -0.39 † |
| Standing long jump (157 – 225 cm) | **-0.59 \*\*** | **-0.41 \*** | **-0.72 \*\*\*** |
| Sit-and-reach (34 – 56 cm) | -0.10 | -0.33 † | -0.23 |
| Whole-body reaction time (222 – 444 msec) | 0.29 | 0.17 | **0.40 \*** |

† p<0.1, \* p<0.05, \*\* p<0.01, \*\*\* p<0.001
Indicate the (minimum value - maximum value) next to the variable name.

The simple correlation analysis revealed a relationship between age in months and performance in all three intervals. Therefore, we examined the correlation between age in months and other factors (Table 2). Consequently, a significant correlation was confirmed with all factors except for body fat mass (range of absolute value, *r* = 0.39 to 0.79; *p* < 0.05). Therefore, age (in months) may have a potential influence on identifying factors that are strongly related to sprint time in each interval.

**Table 2.** Correlation between age in month and other factors.

|  | Age in month |
| --- | --- |
| 0-2.5 m | **-0.49 \*** |
| 2.5-5 m | **-0.47 \*** |
| 5-10 m | **-0.54 \*\*** |
| Height | **0.79 \*\*\*** |
| Weight | **0.53 \*\*** |
| Fat free mass | **0.78 \*\*\*** |
| Body fat mass | 0.03 |
| Trunk rotation (right) | **0.39 \*** |
| Hip flexion angle (left) | **0.44 \*** |
| Hip flexion angle (right) | **-0.69 \*\*\*** |
| Hip flexion angle (left) | **-0.44 \*** |
| Isometric knee extension | **0.69 \*\*\*** |
| Isometric knee flexion | **0.78 \*\*\*** |
| Isotonic knee extension | **0.63 \*\*\*** |
| Isotonic knee flexion | **0.74 \*\*\*** |
| Repeated side-step | **0.67 \*\*\*** |
| Sit-ups | **0.57 \*\*** |
| Standing long jump | **0.78 \*\*\*** |
| Sit-and-reach | **0.44 \*** |
| Whole-body reaction time | **-0.77 \*\*\*** |

\* $p < 0.05$ , \*\* $p < 0.01$ , \*\*\* $p < 0.001$

We conducted partial correlation analyses between each time interval (0‒2.5 m, 2.5-5 m, 5‒ 10 m) and various variables using confirmed age in months with relationships with many variables as a control variable (Table 3). As a result, in the 0‒2.5 m interval, a tendency was observed for faster times with higher hip flexibility (right: *r* = -0.42, *p* = 0.035; left: *r* = -0.60, *p* = 0.001). In the 2.5‒5 m interval, a tendency for faster times with higher trunk flexibility was observed (right: *r* = -0.34, *p* = 0.091; left: *r* = -0.40, *p* = 0.046). In the 5-10m interval, a tendency for faster times with better standing long jump records was observed (*r* = -0.56, *p* = 0.003). Furthermore, the previously non-significant body fat mass in the simple correlation analysis was found to have partial correlations with all interval times (0‒2.5 m: *r* = 0.40, *p* = 0.047; 2.5 m: *r* = 0.37, *p* = 0.071; 5‒10 m: *r* = 0.55, *p* = 0.004).

**Table 3.** Partial correlation coefficients with age in month controlled.

|  | 0-2.5m | 2.5-5m | 5-10m |
| --- | --- | --- | --- |
| 0-2.5m | - | -0.08 | **0.52 \*\*** |
| 2.5-5m | -0.08 | - | -0.03 |
| 5-10m | **0.52 \*\*** | -0.03 | - |
| Height | 0.29 | 0.30 | -0.03 |
| Weight | 0.33 | **0.41 \*** | 0.36 † |
| Fat free mass | 0.14 | 0.37 † | 0.01 |
| Body fat mass | **0.40 \*** | 0.37 † | **0.55 \*\*** |
| Trunk rotation (right) | 0.30 | -0.34 † | 0.17 |
| Trunk rotation (left) | 0.05 | **-0.40 \*** | 0.03 |
| Hip flexion angle (right) | **-0.42 \*** | -0.09 | -0.12 |
| Hip flexion angle (left) | **-0.60 \*\*** | -0.10 | -0.35 † |
| Isometric knee extension | 0.12 | -0.19 | 0.09 |
| Isometric knee flexion | -0.17 | 0.03 | -0.34 † |
| Isotonic knee extension | 0.03 | 0.09 | 0.00 |
| Isotonic knee flexion | 0.24 | -0.03 | -0.07 |
| Repeated side-step | -0.07 | -0.05 | -0.04 |
| Sit-ups | -0.13 | -0.02 | -0.11 |
| Standing long jump | -0.39 † | -0.08 | **-0.56 \*\*** |
| Sit-and-reach | 0.14 | -0.16 | 0.02 |
| Whole-body reaction time | -0.17 | -0.34 | -0.04 |

† p<0.1, \* p<0.05, \*\* p<0.01, \*\*\* p<0.001

## Discussion

This study aimed to identify the key factors that contribute to short sprint performance among junior students. The results of this study showed that in the simple correlation analysis, factors related to the muscular system, such as knee flexion and extension, were confirmed at many intervals. However, in the partial correlation analysis considering the influence of age in months, flexibility factors were strongly related in the first half distance of 10 m, specifically 0‒2.5 m and 2.5‒5 m. When examining each interval, it was confirmed that in the 0‒2.5 m distance, high flexibility of the hip joint, which did not show a relationship in the simple correlation analysis, led to faster times in that distance (pseudo-uncorrelation). In the 2.5‒5 m distance, it was revealed that even when considering the influence of age in months, the flexibility of the trunk, which showed a relationship in the simple correlation analysis, was related. Thus, when considering the influence of age in month, it became clear that especially in the first half segments of 10m, 0‒2.5 m and 2.5‒5 m, sprint performance improves with higher flexibility. In the 5‒10 m distance, the standing long jump, which showed a relationship in both simple and partial correlation analyses, suggests that the ability to generate a horizontal propulsive force is important. Therefore, in the junior period, when considering the influence of age in months, it became evident that factors with strong relationships change even at short distances such as 10 m, hip joint flexibility at 0‒2.5 m, trunk flexibility at 2.5‒5 m, and horizontal propulsive force at 5‒10 m.

In terms of physical elements, body fat mass at all intervals negatively impacted sprint time, even during the junior period. This partly supports previous studies reporting that heavier weights lead to slower sprint times (28). In a previous study that examined body composition and short sprints in female athletes, an increase in body fat percentage led to a shorter first step (1). A longer step length is an important factor for achieving greater acceleration, and an increase in body fat mass affects motion, resulting in slower times. Based on the results obtained from previous studies and the present study, it is clear that an increase in body fat mass leads to an increase in weight, negatively affecting short sprints, even among juniors.

In the initial sections of 0‒2.5 m and 2.5‒5 m, faster times were associated with greater flexibility, enabling longer strides. In this study, it became evident that higher flexibility in the hip joints leads to faster times in the 0‒2.5 m section, while higher flexibility in the trunk results in faster times in the 2.5‒5 m section, highlighting the importance of body flexibility immediately after the start. Previous studies have reported that a greater step width in the 0‒ 5 m section leads to higher acceleration (23). Furthermore, a larger step length in the first step after the start results in greater propulsive force, suggesting immediate performance improvement (30). Taken together, these results suggest that increasing the step length is crucial for achieving high acceleration, with hip and trunk flexibility enabling such a step length.

In particular, hip flexibility is considered specific to the junior period. Previous studies examining power output by joints at the start have reported that hip power output plays a significant role in young athletes, whereas knee power output accounts for a larger proportion (10). Furthermore, high-performing adult top athletes experience more knee flexion at the start (5). Therefore, while the knee plays a crucial role in adult athletes, during the junior period, hip flexibility appears to contribute to the generation of greater power and is closely linked to short sprint performance.

The discovery that trunk flexibility is important at the 2.5‒5 m distance is a novel finding of this study. The intermediate range of 2.5‒5 m showed no relationship with the other time intervals in either the simple correlation analysis or partial correlation analysis, controlling for age in months. This suggests that the 2.5‒5 m range may have independent characteristics. In contrast, correlations were found between the 0‒2.5 m and 5‒10 m intervals. In these two intervals, the 0‒2.5 m range showed a relationship with hip flexibility, while the 5‒10 m range showed a relationship with lower limb function in standing long jump, indicating a commonality in obtaining horizontal propulsive force. Previous studies on short sprints focused on lower limb strength, flexibility, and kinematics (5,7,16). This study focused on upper-body function and found a relationship between upper-body function and performance in the 2.5‒5 m segment of short sprints. Therefore, to improve short sprint performance, which is strongly associated with lower-limb function, the importance of upper-body function and the need to address it as a whole-body movement have been suggested.

In the 5‒10 m distance, the ability to generate horizontal propulsion appeared to contribute to shorter times. Adult athletes exhibit greater horizontal propulsion than junior athletes (17), and a greater horizontal force leads to better sprint performance (6). The standing long jump measured in this study assessed the ability to jump horizontally, and athletes who can jump further excel in generating greater horizontal force. The 5‒10 m distance is regarded as the stage that follows the transition from a stationary start to the running phase and is expected to mark the beginning of the acceleration phase. Therefore, the ability to generate horizontal force may significantly influence performance in this interval.

## Limitations

A limitation of this study is that the step length could not be measured. Previous studies have shown that a larger step length results in greater acceleration; however, this study did not address the relationship between each factor and step length. Therefore, by conducting a three-dimensional motion analysis, it is possible to measure the step length and kinematics of lower limb movements, enabling further detailed analysis.

## Conclusion

This study aimed to investigate the relationship between age, body composition, flexibility, muscle strength, and physical fitness tests during short sprints in junior athletes, as well as the changes in relationships within intervals of 0‒2.5 m, 2.5‒5 m, and 5‒10 m. The results suggested that higher hip flexibility led to faster times in the 0‒2.5 m, higher trunk flexibility led to faster times in the 2.5‒5 m, and better standing long jump performance led to faster times in the 5‒10 m interval. Even at a short distance of 10 m, this study implied that lower body and upper body flexibility were important for the 0‒5 m distance, whereas muscle strength for propulsion became crucial beyond 5 m. The impact of hip flexibility was considered specific to junior athletes, and a novel finding highlighted the contribution of trunk flexibility to sprinting, which often focuses on lower-body function. An increase in body fat mass was suggested to slow down time across all distances. These results underscore the importance of overall flexibility, propulsion force, and body fat regulation in junior short sprints and emphasize the significance of daily training tailored to the sprint distances required in competition.

